# Epigenetic modifiers to treat retinal degenerative diseases

**DOI:** 10.1101/2025.05.22.655558

**Authors:** Evgenya Y. Popova, Lisa Schneper, Aswathy Sebastian, Istvan Albert, Joyce Tombran-Tink, Colin J. Barnstable

## Abstract

We have previously demonstrated the ability of inhibitors of LSD1 and HDAC1 to block rod degeneration, preserve vision, maintain rod-specific transcripts and downregulate those involved in inflammation, gliosis, and cell death in the rd10 mouse model of Retinitis Pigmentosa (RP). To extend our findings we tested the hypothesis that this effect was due to altered chromatin structure by using a range of inhibitors of chromatin condensation to prevent photoreceptor degeneration in the rd10 mouse model. We used inhibitors for G9A/GLP that catalyzes methylation of H3K9, for EZH2 that catalyzes trimethylation of H3K27, and compared them to the actions of inhibitors of LSD1 and HDAC. All the inhibitors decondense chromatin and all preserve, to different extents, retinas from degeneration in rd10 mice, but they act through different metabolic pathways. One group of inhibitors, modifiers for LSD1 and EZH2, demonstrate a high level of maintenance of rod-specific transcripts, activation of Ca^+2^ and Wnt signaling pathways with inhibition of antigen processing and presentation, immune response and microglia phagocytosis. Another group of inhibitors, modifiers for HDAC and G9A/GLP work through upregulation of NGF-stimulated transcription, while down-regulating genes belong to immune response, extracellular matrix, cholesterol signaling and programmed cell death. Our results provide robust support for our hypothesis that inhibition of chromatin condensation can be sufficient to prevent rod death in rd10 mice.

## Introduction

The epigenetic landscape of a cell plays a major role in controlling patterns of gene expression rather than being a passive response to other mechanisms. All cell nuclei contain both euchromatin, which is relatively open and can be actively transcribed, and heterochromatin, where the nucleosomes are more densely packed and transcription is minimal. The transition between these chromatin states is dynamic and under multiple forms of control.

Cells with open euchromatin organization can more easily survive changes in cellular homeostasis in response to stress, but such cells are also prone to cancerous transformation. Cells with more heterochromatic nuclear organization are less susceptible to malignancy but have less ability to adapt to changing environments which makes them predisposed to degeneration and cell death. Mature rod photoreceptors, like most neurons, belong to the second group of cells and have a uniquely closed chromatin organizational structure [1–3]. The restricted transcription potential of such metabolically active cells may increase efficiency under normal conditions but may make them less able to respond under conditions of stress.

Among the retinal degenerations that cause severe vision loss, Retinitis Pigmentosa (RP) affects 1.5 million people worldwide. It is inherited in a Mendelian fashion and different forms can be manifested in children and adults. RP is characterized by a progressive loss of rod photoreceptors followed by a secondary deterioration of cone photoreceptors, the latter making the disease particularly debilitating [4]. RP is heterogeneous with >4000 identified mutations in >300 genes/loci [5]. The link between mutant gene and cell death is not always clear. Some mutant proteins are toxic and directly induce cell death while others perturb metabolic networks that indirectly lead to cell death. The large numbers of different mutations and pathways leading to cell death have hindered identification of suitable RP treatments. There is currently no cure for RP, although gene therapy, prosthetic implants and pharmacological agents are all being explored as therapeutic approaches [6–8]. The most successful approaches for RP therapy are likely to be gene-independent pharmacological advances with potential to target multiple forms of the disease and which can be used for early stage treatment.

Epigenetic remodeling of gene expression is a powerful new strategy to fight neurodegeneration. Degenerative diseases of the retina, including RP, are good candidates for this approach because there are both excellent animal models and an available patient population, but no treatments. In our previous published work [9] we demonstrated that inhibition of LSD1and HDAC1 in a mouse model of RP altered chromatin structure and led to preservation of rod photoreceptors and visual function, retaining expression of rod-specific genes, with decreased inflammation, cell death and Muller cell gliosis. Because LSD1 and HDAC1 themselves can affect inflammation through effects of NFkB activation [10–12], these results could not fully resolve the importance of chromatin remodeling.

To address this issue, we have now broadened our studies to test whether multiple classes of inhibitors of chromatin condensation can prevent rod death. We have chosen two enzymes of chromatin organization that act differently to those we have used previously. The first enzyme, Enhancer of zeste homolog 2 (EZH2), is a histone lysine-N-methyltransferase that catalyzes trimethylation of H3K27. It is a component of the Polycomb Repressive Complex 2 (PRC2) and is critical for both embryonic development and gene silencing. Its methylation activity facilitates both the formation of heterochromatin and heterochromatin remodeling.

DZNep (3-deazneplanocin A) and UNC1999 are inhibitors of EZH2 that selectively blocks formation of H3K27me3.Treatment with DZNep has been reported to slightly delay photoreceptor degeneration in rd1 mice, and cause decreased microglial activation in models of ischemic stroke [13, 14] and UNC1999 intravitreal injection partly prevent retina degeneration in two mouse models [15].

The second enzyme, G9A/GLP, is a heterodimeric complex that catalyzes mono- and di- methylation of H3K9. The inhibitor UNC0642 is a potent and highly selective inhibitor of this complex and improved cognition and reduced oxidative stress and neuroinflammation in a mouse model of Alzheimer disease[16]. It also improved survival in mouse model of Prafer-Willi syndrome [17].

Our results demonstrated that all these inhibitors that decondense chromatin decrease retinal degeneration in mouse model of RP, rd10, but they act through different metabolic pathways. Based on our results we propose that these epigenetic inhibitors cause more open and accessible chromatin in rod photoreceptors, allowing expression of clusters of neuroprotective genes. A second mechanism that also promotes rod photoreceptor survival is an action of epigenetic inhibitors leading to suppression of inflammatory processes in microglia and glia.

## Methods

### Antibodies and reagents

Chemicals were purchased from Fisher Scientific (Pittsburgh, PA), unless otherwise noted. 0.9% bacteriostatic sodium chloride was from APP Pharmaceuticals (Schaumburg, IL). LSD1 inhibitor GSK2879552, EZH2 inhibitors DZNep and UNC1999 was from Selleckchem.com (Huston, TX). The HDAC inhibitor Romidepsin and G9A/GLP inhibitor UNC0642 were from Sigma (St. Louis, MO), anti-rhodopsin (RHO) monoclonal antibodies have been described previously [18] and react with an N-terminal sequence shared by many species. The commercial antibodies used were: anti-GFAP (MAB360, Millipore, Temecula, CA), anti-IBA1 (for the Aif1 gene; 019-19741, Wako, Richmond, VA).

### Animals

Wild type C57Bl/6J (cat # 000664), and rd10 B6.CXB1-Pde6brd10/Jrd10 (cat#004297) mice were purchased from Jackson laboratory (Bar Harbor, ME, United States) and housed in a room with an ambient temperature of 25°C, 30–70% humidity, a 12-h light–dark cycle, and ad libitum access to rodent chow. This study was carried out using both male and female mice in accordance with the National Research Council’s Guide for the Care and Use of Laboratory Animals (8th edition) and all animal experiments were approved by the Pennsylvania State University College of Medicine Institutional Animal Care and Use Committee (protocol #46993).

To cell sorting of rod photoreceptors expressing GFP from degenerated and GSK treated retina we crossed rd10 mouse (B6.CVB1-Pde6brd10/j) with mouse B6.Cg-Tg(Nrl-EGFP)1Asw/J (gift from Dr. A. Swaroop) [19] to establish mouse line that harbors both homozygous rd10 mutation and homozygous expression of green fluorescence protein in rod photoreceptors.

Genotyping was done according to Jackson lab protocols for rd10 mice (cat #004297) and B6.Cg-Tg(Nrl-EGFP)1Asw/J (cat#021232).

### Treatment with inhibitors

All inhibitors were diluted in specific diluent or in 0.9% bacteriostatic sodium chloride (saline). Mice were treated daily with intraperitoneal injections (i.p.) of UNC1999 at 15mg/kg (diluted in10% DMSO,40% PEG300, 5% Tween-80 and 45% saline), GSK2879552 (GSK, diluted in saline) at 4.2mg/kg, DZNep at 1-1.5 mg/kg (diluted in saline), Romidepsin at 0.2mg/kg (diluted in saline), UNC0642 at 7.5mg/kg (diluted in saline) or saline(diluent) as control.

### Tissue collection

Whole retinas were isolated from animals by removing the sclera and most of the retinal pigmented epithelium (RPE) layer under PBS. Right eye retinas from each animal were taken for RNA extraction, cDNA preparation and RT-PCR. Immediately after isolation, tissue was flash frozen in liquid nitrogen and stored at -80°C. Left eye retinas from each animal were subjected to fixation, cryopreservation, sectioning and immunofluorescence staining.

### FACS

We treated mice from this newly established line from PN9 till PN24 with daily i.p. injections of saline (control) or GSK2879552 at 4.2mg/kg. Mouse retinas were dissected and dissociate according to a protocol from Solovei’s lab [20]. Cells were sorted on BD FACSDiva 9.0.1 in Flow Cytometry Core at Penn State College of Medicine. RNA was isolated from 2-3 x 10^6^ sorted cells per sample with RNeasy Mini Kit and RNA shredder (Qiagen). RNA integrity number (RIN) was measured using BioAnalyzer (Agilent Technologies) RNA 6000 Nano Kit to confirm RIN above 9 for each sample.

### RNA extraction and cDNA preparation

RNA extraction and purification followed the manufacturer’s protocol from RNeasy Mini Kit and RNA shredder (Qiagen). Final RNA concentrations were determined spectrophotometrically using a NanoDrop 1000 Spectrophotometer (Thermo Fisher Scientific, Wilmington, Delaware). cDNA was synthesized with SuperScript III First-Strand Synthesis System kit according to manufacturer’s protocol (Invitrogen, Carlsbad, California).

### RT-PCR

Primers were purchased from Integrated DNA Technologies (IDT). The sequence information is listed in Supplemental Table 1. For quantitative real-time PCR we used 2x iQ-SYBR Green PCR supermix from Bio-Rad. Samples in triplicate were run on an iQ5 Multicolor Real Time PCR Detection System (Bio-Rad). The relative expression level for each gene was calculated by the 2-ΔΔCt method and normalized to GAPDH. Genes were considered up or down-regulated if p value <0.05.

### RNA-seq

RNA was extracted from PN24 retina rd10 animals treated with saline or epigenetic inhibitors from PN9 till PN24. RNA integrity number (RIN) was measured using BioAnalyzer (Agilent Technologies) RNA 6000 Nano Kit to confirm RIN above 7 for each sample. RNA-seq libraries were prepared in the Penn State College of Medicine Genome Sciences core (RRID:SCR_021123) using the Stranded mRNA Prep, Ligation kit (Illumina) as per the manufacturer’s protocol. Briefly, polyA RNA was purified from 200 ng of total RNA using oligo (dT) beads. The extracted mRNA fraction was subjected to fragmentation, reverse transcription, end repair, 3’ - end adenylation, and adaptor ligation, followed by PCR amplification and magnetic bead purification (Omega Bio-Tek). The unique dual index sequences (IDT® for Illumina RNA UD Indexes Set B, Ligation, Illumina) were incorporated in the adaptors for multiplexed high-throughput sequencing. The final product was assessed for its size distribution and concentration using BioAnalyzer High Sensitivity DNA Kit (Agilent Technologies). The libraries were pooled and sequenced on Illumina NovaSeq 6000 (Illumina), to get an average of 25 million, paired end 50 bp reads, according to the manufacturer’s instructions. Samples were demultiplexed using bclconvert software (Illlumina). Adaptors were not trimmed during demultiplexing.

BBDuk (version 38.69) was used to trim adapters and filter low-quality sequences using “qtrim=lr trimq=10 maq=10” options and the adapter database provided. Next, alignment of the filtered reads to the mouse reference genome (mouse Ensembl release 67 (GRCm37 / NCBIM37 / mm9)) was done using HISAT2 [21] (version 2.2.1) applying --no-mixed and --no-discordant options. Read counts were calculated using HTSeq [22] (version 0.11.2) by supplementing Ensembl gene annotation (release 67: "Mus_musculus.Ensembl.NCBIM37.67.gtf"). Processing of the read counts and differential gene expression were performed with the EdgeR R package [23] (versions 3.42.4 and 4.3.1, respectively). Genes with low counts were filtered using the filterByExpr function. After removal, library sizes were normalized using the calcNormFactors function. The variability in gene expression across all samples was accounted for by estimating the dispersion for each gene using the estimateDisp function. Negative Binomial Generalized Linear Models (NB GLMs) were the fitted to the count data using the glmFit function. Differentially expressed genes were then determined by the likelihood ratio test method (glmLRT function). Significance was defined to be those with q-value < 0.05 calculated by the Benjamini-Hochberg method to control the false discovery rate (FDR) and log2 fold change is greater than 1 or smaller than -1. The pheatmap R package was used for generating heatmaps. Raw counts and differential expression analysis generated during this study are available at GEO Submission **GSE295538**.

### RNA-seq for FACS-sorted rod photoreceptors.

RNA-seq libraries were prepared in the Penn State College of Medicine Genome Sciences core (RRID:SCR_021123) using the SMART-Seq mRNA HT LP kit (Takara Bio) as per the manufacturer’s instructions. Briefly, cDNA was synthesized from 1 ng of RNA with recommended PCR cycles of 11 and purified with magnetic beads. Quality and quantity of the cDNA was checked with BioAnalyzer High Sensitivity DNA Kit (Agilent Technologies). Sequencing libraries were then prepared with 300 pg of the cDNA and indexed with SMARTer RNA Unique Dual Index Kit. The final product was assessed for its size distribution and concentration using BioAnalyzer High Sensitivity DNA Kit. The libraries were pooled and sequenced on Illumina NovaSeq 6000 (Illumina), to get on average 25 million, paired end 50 bp reads, according to the manufacturer’s instructions.

Samples were demultiplexed using bclconvert software (Illlumina). Adaptors were not trimmed during demultiplexing.

### Pathway, gene ontology and upstream regulator analysis

The clusterProfiler R package [24] was used to perform GO functional analysis an plot the output. GO terms were reduced using the rrvgo R package and p-values used for plotting were the least significant. Ingenuity Pathways Analysis (IPA) was used to identify upstream regulators and significantly enriched canonical pathways with following cutoff of FDR <0.05 and fold change in gene expression bigger than 2 or smaller than 0.5.

### Immunofluorescence Staining

Retinas were fixed in 4% paraformaldehyde overnight at 4°C, washed in PBS, incubated in 5% sucrose/PBS for 30 min and then cryopreserved in 20% sucrose/PBS overnight at 4°C. Retinas were embedded in 2:1 mix of 20% sucrose and OCT (Sakura Finetek Torrance, CA) and stored at -80°C. Blocks with tissue samples were sectioned to 7-10µm on a Cryostat Microtome HM550 (Thermo Fisher Scientific) and stored at - 20°C. Antigen retrieval was performed by incubating the slides in 10mM sodium citrate pH 6 for 30 min at 80°C. Double labeling immunohistochemistry was performed as previously described [25, 26] using fluorescent Alexa Fluor-conjugated secondary antibodies diluted 1:800 (Invitrogen, Carlsbad, California). Primary antibodies were diluted as follow: anti-RHO 1:50, anti-GFAP 1:1000, anti-IBA1 1:450 (Aif1 gene). Slides were counterstained with Hoechst 33258 (1mg/ml diluted 1:1000) and visualized using a Zeiss confocal microscope LSM900. The acquisition parameters were maintained constant for each set of experiments. Fluorescence intensity was assessed using ImageJ software (Bethesda, MD).

### Statistical Analyses

Results are presented as means ± standard error of the mean (SEM). Unpaired, one tail Student’s t-test (two-tailed, unpaired) was used to evaluated statistical significance between groups. P value < 0.05 was considered significant. Statistical analyses for experiments were performed using the GraphPad Prism software.

## Results

### 1. Treatment of rd10 mouse model of RP with EZH2 inhibitors leads to retina neuroprotection

We first treated rd10 mice with EZH2 inhibitor DZNep. Systemic administration of DZNep to mice could be deleterious, so we carried out a series of experiments to investigate the best timetable of DZNep treatment. The optimal drug administration schedule for animal survival was from PN9 to PN12 each day 1.5mg/kg DZNep and after it, from PN13 to PN24 each second day with 1mg/kg DZNep. Immunostaining demonstrated preservation of retina outer nuclear layer of rods (ONL) (Fig.1A) and down regulated of GFAP activation after DZNep treatment. After establishing that treatment of rd10 with inhibitor specific for EZH2 led to preservation of rod photoreceptors we next studied how this treatment influences gene expression. We performed RNA-seq on retina samples from rd10 mice treated with DZNep or saline from PN9 till PN24 (Fig.1B). Using a cutoff of FRD <0.05 and a Log (fold change)>1, rd10 mice treated with this inhibitor have 518 genes upregulated and 367 genes downregulated (Fig.1B). Ingenuity Pathways Analysis (IPA) demonstrated that the Visual Phototransduction pathway was the most upregulated pathway, while most downregulated pathways were related to inflammation and immuno-response – Integrin cell surface interactions and Interferon gamma signaling (Fig.1C). Cnet plots made from Gene Ontology (GO) functional enrichment analysis showed specific genes that are involved in the upregulated pathways hub of visual perception, detection of light and sensory perception of light stimulus, as well as the downregulated genes hub of adaptive immune response and antigen processing and presentation (Fig.1D). Top upstream regulators according to IPA were RHO, Immunoglobulin, CRX, TGFB1, TNF and KDM1a (LSD1). We confirmed RNA-seq results with qRT-PCR for selected genes. Rod- specific genes were upregulated and gene markers for inflammation were down regulated under DZNep inhibition in rd10 mice (Sup. Fig.1A). GO analysis identified that the 10 most down-regulated Molecular Functions (MF) were related to immunity and extracellular matrix, the 10 most down-regulated Cell Components (CC) were connected to plasma membrane and MHC complexes, and the 10 most down-regulated Biological Process (BP) were all linked to immunity (Sup.Fig.1B). GO analysis identified that the 2 most up-regulated Molecular Functions (MF) were retinoid binding and cyclic nucleotide-gated channel activity, the 5 most up-regulated Cell Components (CC) included photoreceptor cilium, plasma membrane and photoreceptor inner segment, and the 10 most up-regulated Biological Process (BP) were all linked to eye development and function (Sup.Fig.1C). Thus, the GO analysis for differently expressed genes (DEG) between control and DZNep treatment group corroborated the IPA results.

**Fig. 1.**
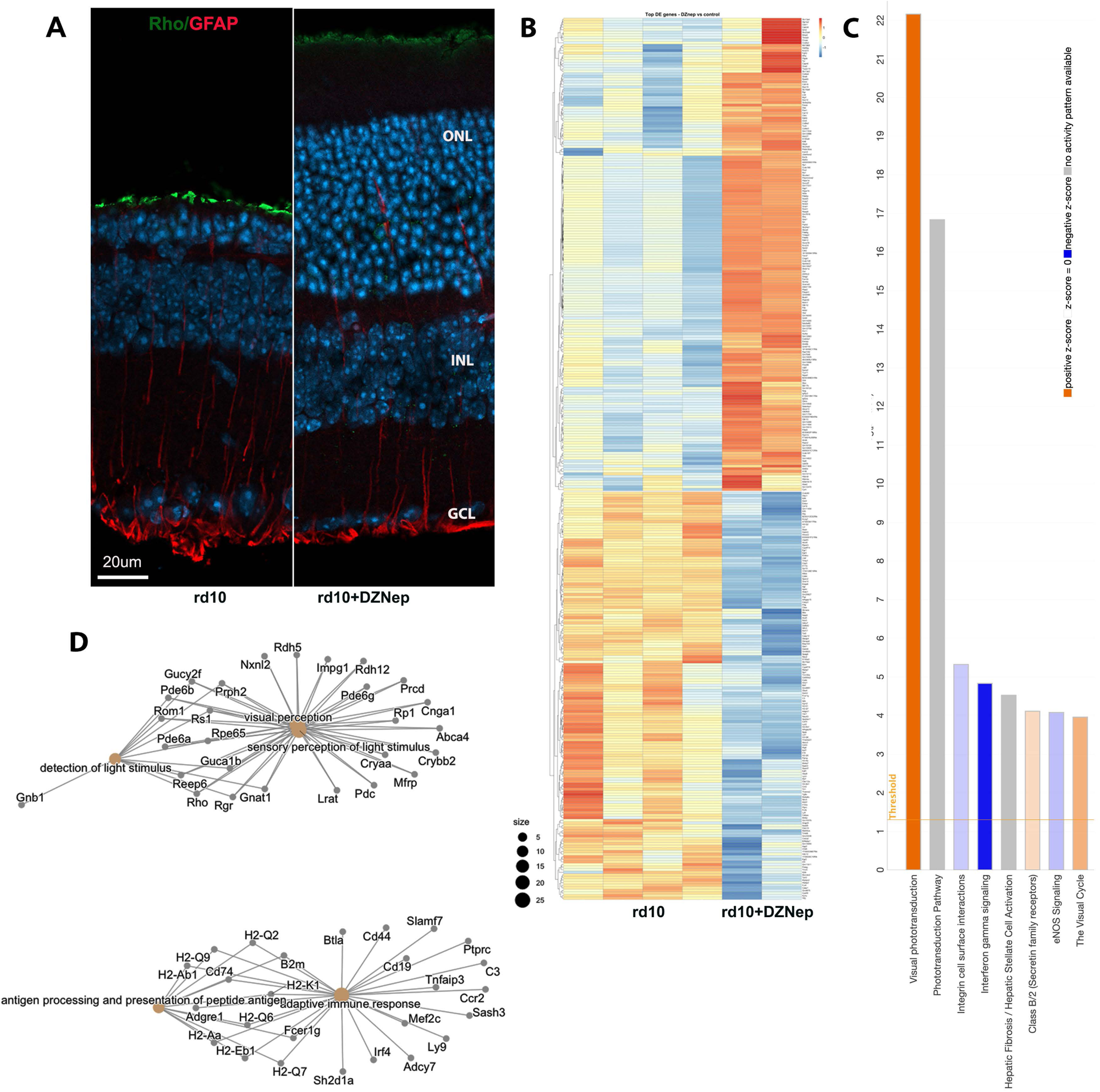
Treatment of rd10 mouse model of retinitis pigmentosa with EZH2 inhibitor DZNep leads to retina neuroprotection. **A**. Immunofluorescence microscopic images of retina sections from PN24 rd10 mice treated from PN9 till PN24 with DZNep or only saline (control), stained with RHO (green), GFAP (red), and nuclear counterstained with Hoechst33358. **B.** RNA-seq analysis of altered retinal gene expression under DZNep inhibition. Heatmap of DEG between rd10 and treated with DZNep rd10 retinas (FDR<0.05; FC - greater than 2 or smaller than 0.5). **C.** Top Ingenuity canonical pathways for DEG between rd10 and treated with DZNep rd10 retinas (FDR<0.05; FC - greater than 2 or smaller than 0.5) according to IPA. **D.** Category-gene network plot depicting DEG between rd10 and treated with DZNep rd10 retinas.

We compared these results with those using a more specific EZH2 inhibitor, UNC1999, to treat the rd10 mouse model. UNC1999 is less toxic for mice and was used at 15 mg/kg from PN9 to PN24. UNC1999 action on rd10 mice was very similar to DZNep, and showed preservation of ONL and down regulation of Muller glia activation (Fig.2A). The number of photoreceptor rows in ONL increased from an average of 5 in rd10 to 10 in retinas of rd10 mice treated with UNC1999 (Fig.2B). We performed RNA-seq on retina samples from rd10 mice treated with UNC1999 or saline from PN9 till PN24 (Fig.2C). rd10 mice treated with this inhibitor have 368 genes upregulated and 519 genes down-regulated (Fig.2C). RNA-seq IPA analysis similarly demonstrated preservation of expression of rod-specific and visual phototransduction pathway genes and down regulation of neuroinflammation genes and complement system pathway (Fig.2D). Top upstream regulators according to IPA were LPS, TNF, RHO, KDM1a (LSD1), IFNG and Immunoglobulin. We confirmed the RNA-seq results with qRT-PCR for selected genes. Rod-specific genes were upregulated and genes specific for inflammation were down regulated under UNC1999 inhibition in rd10 mice (Sup.Fig.2A). GO analysis identified that the 10 most down-regulated Molecular Functions (MF) were related to immunity and transmembrane receptor binding, the 10 most down-regulated Cell Components (CC) were connected to plasma membrane and MHC complexes and the 10 most down- regulated Biological Processes (BP) were all linked to immunity (Sup.Fig.2B). GO analysis identified that the 2 most up-regulated Molecular Functions (MF) were anion transporter activity and extracellular matrix structure, the 10 most up-regulated Cell Components (CC) included photoreceptor cilium, plasma membrane and photoreceptor inner segment, and the 10 most up- regulated Biological Processes (BP) were all linked to eye development and function (Sup.Fig.2C). Thus, the GO analysis for differently expressed genes (DEG) between control and UNC1999 treatment group corroborated the IPA results.

**Fig. 2.**
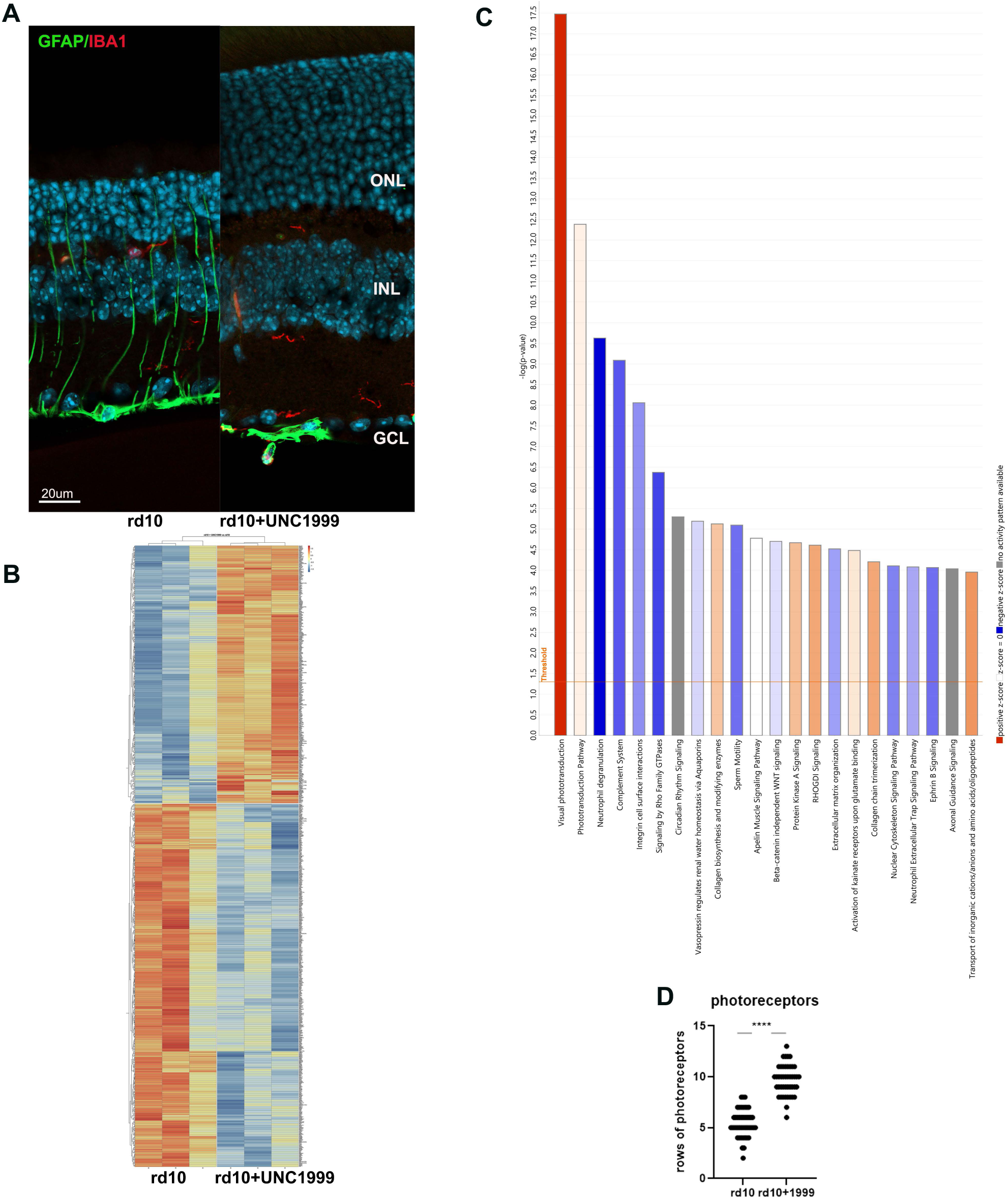
Treatment of rd10 mouse model of retinitis pigmentosa with EZH2 inhibitor UNC1999 leads to retina preservation. **A**. Immunofluorescence microscopic images of retina sections from PN24 rd10 mice treated from PN9 till PN24 with UNC1999 or only saline (control), stained with GFAP (green), IBA1 (red), and nuclear counterstained with Hoechst33358. **B.** RNA-seq analysis of altered retinal gene expression under UNC1999 inhibition. Heatmap of DEG between rd10 and treated with UNC1999 rd10 retinas (FDR<0.05; FC - greater than 2 or smaller than 0.5). **C.** Top Ingenuity canonical pathways for DEG between rd10 and treated with UNC1999 rd10 retinas (FDR<0.05; FC - greater than 2 or smaller than 0.5) according to IPA. **D.** Rods rows were counted in central retina of rd10 PN24 mice treated from PN9 till PN24 with UNC1999 or only with saline for 3 biological and 3 technical replicas (± SEM, **** p<0.0001).

### 2. Treatment of rd10 mouse model of RP with G9A/GLP inhibitor leads to partial retina neuroprotection

During rod photoreceptor maturation their nuclei undergo dramatic transformation, so that in adult retina rods have only one big heterochromatic focus that occupies much of the nuclei volume. The epigenetic mark H3K9me2 is associated with facultative heterochromatin, so we used UNC0642, inhibitor of the heterodimeric complex G9A/GLP that catalyzes mono- and di- methylation of histone H3K9, to study if relaxing heterochromatin in rd10 mice in such way will help rod photoreceptor to survive. Treatment with UNC0642 inhibitor partially preserved rod photoreceptor and retina layer structure and lower gliosis (Fig.3A). The number of photoreceptors rows in ONL increased from an average of 4 for rd10 to 7-8 in retinas of rd10 mice treated with UNC0642 (Fig.3B). The lower efficacy of UNC0642 was probably due to low solubility in aqueous solutions and known low permeability across the blood brain barrier [27]. We performed RNA-seq on retina samples from rd10 mice treated with UNC0642 or saline from PN9 till PN24 (Fig.3C). rd10 mice treated with this inhibitor have only 44 genes upregulated and 28 genes downregulated (Fig.3C). IPA demonstrated that this limited set of genes participate mostly in upregulating stress response, including ERK5 signaling, NGF-stimulated transcription, NRF2-medited oxidative response and down regulation of EIF2 and DHCR24 signaling responses (Fig.3D). EIF2 signaling is connected with upregulation of transcription and DHCR24 signaling - with cholesterol synthesis.

**Fig. 3.**
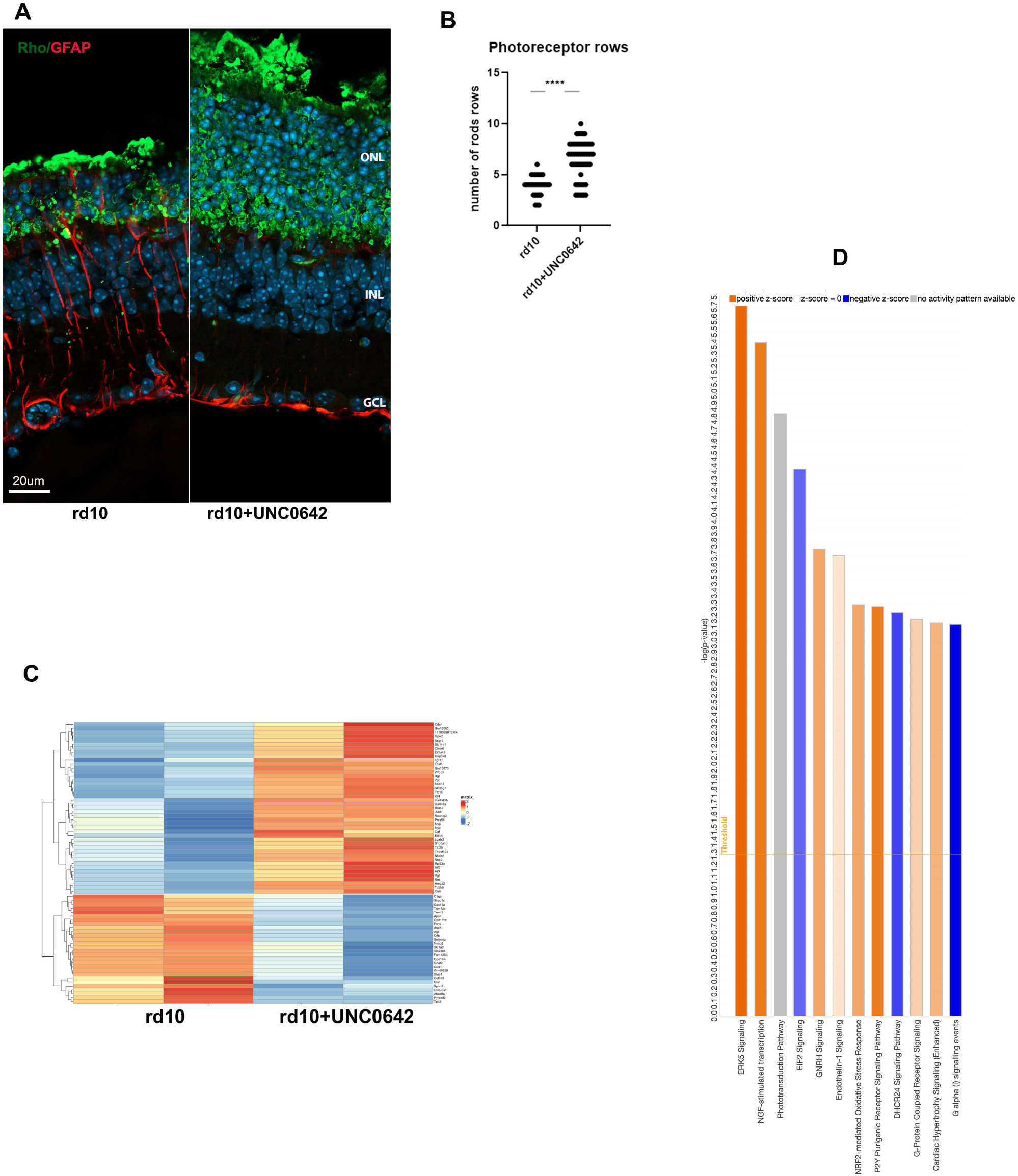
Treatment of rd10 mouse model of retinitis pigmentosa with G9a/GLP inhibitor UNC0642 leads to partial retina protection. **A**. Immunofluorescence microscopic images of retina sections from PN24 rd10 mice treated from PN9 till PN24 with UNC0642 or only saline (control), stained with RHO (green), GFAP (red), and nuclear counterstained with Hoechst33358. **B.** Rods rows were counted in central retina of rd10 PN24 mice treated from PN9 till PN24 with UNC0642 or only with saline for 2-3 biological and 3 technical replicas (± SEM, **** p<0.0001). **C.** RNA-seq analysis of altered retinal gene expression under UNC0642 inhibition. Heatmap of DEG between rd10 and treated with UNC0642 rd10 retinas (FDR<0.05; FC - greater than 2 or smaller than 0.5). **D.** Top Ingenuity canonical pathways for DEG between rd10 and treated with UNC0642 rd10 retinas (FDR<0.05; FC - greater than 2 or smaller than 0.5) according to IPA.

### 3. RNA-seq of retina of rd10 mice treated with HDAC1 inhibitor

In our previous paper [9] we demonstrated that just as inhibitors of LSD1 block rod photoreceptor death in rd10 mice an inhibitor of histone deacetylase HDAC1, Romidepsin has similar effect (see [9]- Fig.1). While under Romidepsin inhibition rod-specific genes were upregulated to much lesser extent as under GSK treatment (see [9]- Fig.5A), down regulation of immune and inflammatory specific genes was robust (see [9]- Fig.5F). For proper comparison between different epigenetic inhibitors we performed RNA-seq on retina samples from rd10 mice treated with Romidepsin or saline from PN9 till PN24 (Fig.4A). rd10 mice treated with this inhibitor have 511 genes upregulated and 402 genes downregulated (Fig.4A). Rod-specific genes such as Rho, Aipl1, Pde6b, Ne2e3, Guca1a, Pdc, Ptp4a3 were upregulated, while cone-specific genes Gnat2, Pde6c and Opn1mw were downregulated. But mostly genes with altered expression belong to immune and inflammation response pathways. IPA demonstrated association of HDAC inhibition with inflammation and extracellular matrix organization pathways (Fig. 4B). Top upstream regulators according to IPA were LPS, b-estradiol, 1L1b, immunoglobulin, IFNG, IL6. Cnet plots made from Gene Ontology (GO) functional enrichment analysis showed specific DEG that are involved in pathways hubs of response to bacteria, immune effector process, regulation of defense response and negative regulation of viral process (Fig.4C). GO analysis identified that the 10 most down- regulated Molecular Function (MF) were related to receptor activity, extracellular matrix and binding to it, the 5 most down- regulated Cell Component (CC) were connected to extracellular matrix and MHC complexes and the 10 most down-regulated Biological Process (BP) were all linked to immunity, defense response and NF-kB signaling (Sup.Fig.4A). GO analysis identified that the 2 most up-regulated Molecular Function (MF) were DNA-binding/ RNA-polymerase transcription and cyclin-dependent inhibitor activity and the 10 most up-regulated Biological Process (BP) were all linked to cell differentiation and proliferation (Sup.Fig.4B). Thus, GO analysis for DEG between control and Romidepsin treatment group corroborated with IPA.

**Fig. 4.**
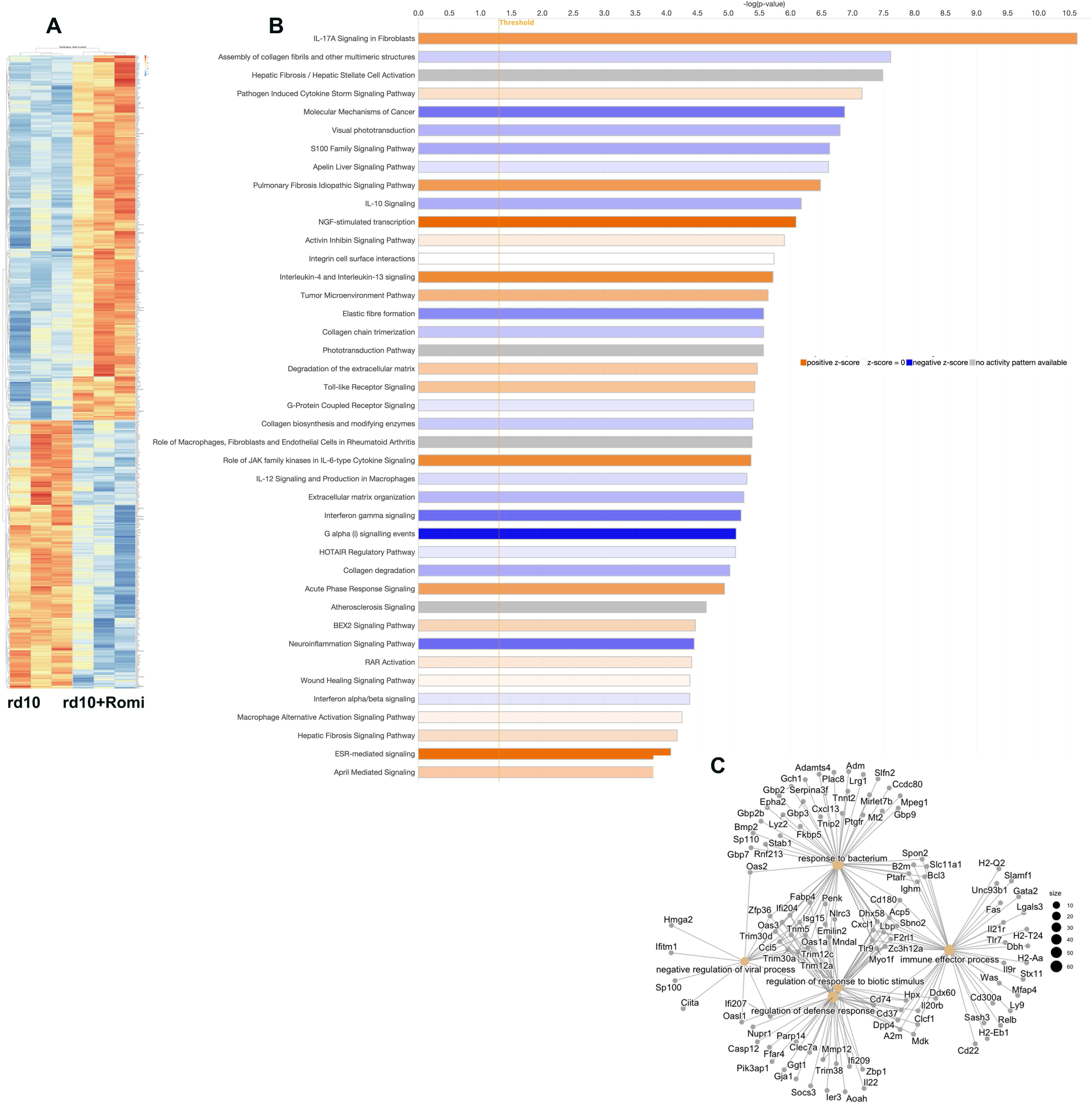
RNA-seq analysis of altered retinal gene expression in rd10 mouse model of retinitis pigmentosa under HDAC inhibition with Romidepsin. **A.** Heatmap of DEG between rd10 and treated with Romidepsin rd10 retinas (FDR<0.05; FC - greater than 2 or smaller than 0.5). **B.** Top Ingenuity canonical pathways for DEG between rd10 and treated with Romidepsin rd10 retinas (FDR<0.05; FC - greater than 2 or smaller than 0.5) according to IPA. **D.** Network plot of enrichment functions by cnetplots for DEG between rd10 and treated with Romidepsin rd10 retinas.

**Fig. 5.**
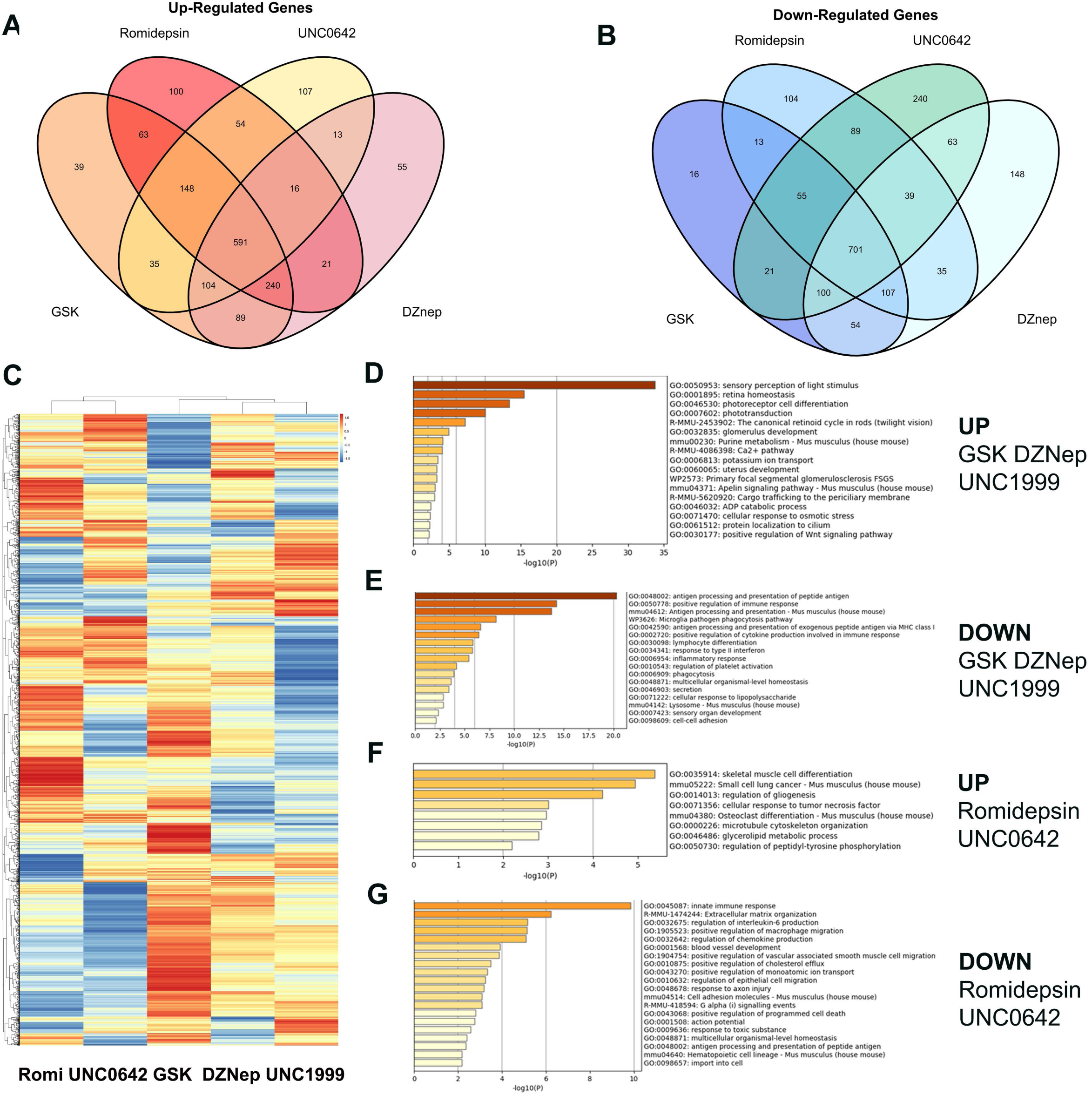
Comparison of action of different inhibitors of chromatin compaction on gene expression in rd10 mouse model of retinitis pigmentosa. **A.** Venn diagram representing overlapping upregulated DEG identified by RNA-seq analysis among GSK, DZNep, UBC0642 and Romidepsin treated groups. **B.** Venn diagram representing overlapping down regulated DEG identified by RNA-seq analysis among GSK, DZNep, UBC0642 and Romidepsin treated groups. **C.** Heatmap for comparison of DEG identified by RNA-seq analysis among GSK, DZNep, UBC0642, UNC1999 and Romidepsin treated groups. **D – G.** Barplots representing enriched ontology clusters for common upregulated (**D**) and downregulated (**E**) DEG identified by RNA-seq analysis among GSK, DZNep and UNC1999 treated groups; and for common upregulated (**F**) and downregulated (**G**) DEG identified by RNA-seq analysis among Romidepsin and UNC0642 treated groups.

### 4. Comparison of action of different inhibitors of chromatin compaction

All of the inhibitors of chromatin compaction: Romidepsin for HDAC, DZNep and UNC1999 for EZH2 and G9A/GLP inhibitor UNC0642 preserved rod photoreceptors from degeneration in the mouse model of RP, rd10. To find common biological pathways and mechanisms to fight retina degeneration we compare mRNA-seq data for samples presented in this manuscript and related it to our previous mRNA-seq data for GSK2879552 treated retinas [9]. The overall comparison revealed that one group of epigenetic inhibitors: DZNep, UNC1999 and GSK have much more common upregulated genes then another group, UNC0642 and Romidepsin (Fig.5A, B).

Approximately half of the common upregulated DEGs for GSK, DZNep and UNC1999 treatments belong to visual phototransduction and visual cycle pathways and included such retina developmental TFs as Nrl and Cazs1 (Fig.5D). In addition, there were several common up-regulated pathways that are important to support photoreceptor survival: Ca^+2^ metabolism, potassium ion transport and positive regulation of Wnt signaling pathway (Fig.5D). Common down-regulated DEGs for GSK, DZNep and UNC1999 treatments belong to cellular immune response pathways, antigen processing and presentation, and lymphocyte differentiation (Fig.5E).

Treatments with Romidepsin and UNC0642 gave different results from the other three inhibitors. Fewer genes belonging to phototransduction and visual cycle were robustly upregulated in this group, though a number of others were upregulated to a lesser extent. Several of upregulated genes are members of Immediate early response genes (IER) – Fosb, Fosl1, Gadd45a, Gadd45b, Myc, Junb. These genes participate in NGF-stimulated transcription, transcriptional activation and NRF2-mediated oxidative stress responses. Common up- regulated processes for these inhibitors include cell differentiation, cellular response to TNFa, cytoskeleton organization (Fig.5F). Common down-regulated DEGs for Romidepsin and UNC0642 treatments belong to pathways associated with extracellular matrix, neuroinflammation signaling and cholesterol and programmed cell death (Fig.5G).

### 5. Changes in gene expression of isolated rod photoreceptor in rd10 mice treated with GSK2879552

Whole retina RNA-seq gives a complex picture of gene expression changes especially for low expressing transcript as different types of cell could demonstrate opposite changes for the same gene. Even for the largest cell population such as rod photoreceptors changes in low-expressed but important for rod genes could be skewed because of altered expression of the same gene in other cell types. To identify changes specific for rod photoreceptors we crossed rd10 mice with mice that expressed EGFP in rods under the Nrl promoter [19] and established a mouse line that is homozygous for both the rd10 mutation and expression of green fluorescence protein in rod photoreceptors. Treatment of this line with GSK had same protective effect as treatment of rd10 (data not shown). Pure rod photoreceptors were FACS sorted from dissociated retinas of saline- and GSK-treated animals and rod specific RNA was subjected to RNA-seq. Sorted rod photoreceptors had 25 genes that were down-regulated and 46 genes that were up regulated (Fig.6A). Among the down-regulated genes were Edn2, a known marker of retina degeneration, Gadd45b, a stress response gene, and Igrm1, a gene that positively regulates macroautophagy. Two cone specific genes, Pde6h and Gnb3 were also down regulated. The majority of upregulated genes had low expression at PN24 (according to [28]; Visualization Community | St. Jude Cloud), 7 genes are located in retina developmental enhancer, cis-regulatory DNA sequences that orchestrating gene expression during retina development. One third of upregulated genes resided in heterochromatin, again emphasizing that treatment with epigenetic inhibitors decondensed heterochromatin, leading to re-expression of these genes. Two examples of genes residing in rod photoreceptor heterochromatin and having no retina expression in at this age, but became upregulated in retinas treated with GSK: *Kit*, important for regeneration and homeostasis; and *Sstr*, important for photoreceptor development and inhibits calcium entry by suppressing voltage-dependent calcium channels.

**Fig. 6.**
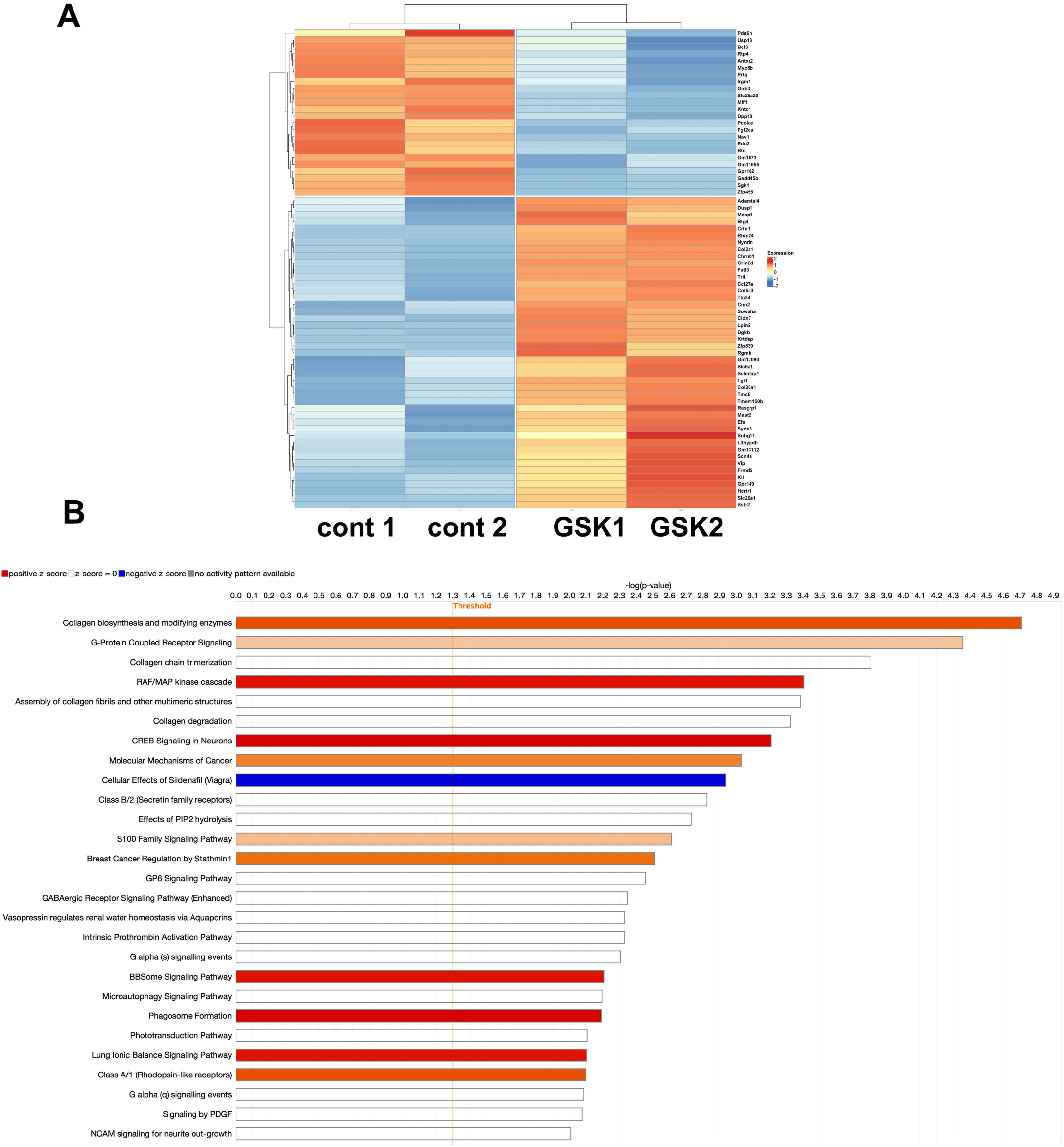
RNA-seq analysis of altered gene expression of FACS-sorted rods from rd10 mouse model of retinitis pigmentosa treated with GSK2879552 vs untreated rd10. **A.** Heatmap of DEG between rd10 and treated with GSK rd10 rod photoreceptors (FDR<0.05; FC - greater than 2 or smaller than 0.5). **B.** Top Ingenuity canonical pathways for DEG between rd10 and treated with GSK rd10 rod photoreceptors (FDR<0.05; FC - greater than 2 or smaller than 0.5) according to IPA.

According to IPA (Fig.6B) most upregulated pathways by GSK in the sorted rods were related to collagen metabolism, G-protein coupled receptor signaling, RAF/Map kinase cascade and CREB-signaling in neurons.

We compared our RNA-seq data on sorted rods (rd10 vs rd10+ GSK) and data of rod photoreceptors from scRNA-seq of rd10 mice at PN21 (cluster C1 vs clusters C2, C3) [29]. We used the same relaxed cutoff for our data that was used in the scRNA-seq: p<0.05, logFC>0.25. There were 41 genes in common between the two sets. Most of these genes were upregulated during degeneration of rods and downregulated to near normal levels under GSK treatment.

These included genes regulating apoptosis or stress response genes *Cd47, E2f6, Gadd45b, Edn2, Wwtr1*. Interestingly, there were only 3 genes, that downregulated during degeneration but upregulated under GSK treatment. They are: 1).*HK2*, hexokinase, that functions in negative regulation of mitochondrial membrane permeability, negative regulation of reactive oxygen species metabolism, anti-apoptotic, photoreceptor protection; 2). *Rabgef1*, with functions in ubiquitin protein ligase activity, dendritic transport, loss of it causes aberrant morphogenesis and altered autophagy in photoreceptors leading to retinal degeneration; 3). *Selenbp1*- selenium binding protein 1, that catalyzes the oxidation of methanethiol and plays a role in intra-Golgi protein transport. In addition to the effects of GSK in sorted rod photoreceptor, all three were upregulated in rd10 treated with DZNep, Romidepsin und UNC1999.

## Discussion

In our previous study we found that epigenetic inhibitors GSK2879552 and Romidepsin block rod degeneration, preserve vision, maintain rod-specific transcripts and downregulate those involved in inflammation, gliosis, and cell death in a mouse model of RP, rd10 mice [9]. GSK2879552 and Romidepsin inhibit different enzymes, LSD1 and HDACs respectively, but these enzymes have common characteristics - they condense chromatin inside nucleus of the cell and sometime work in the same protein complex.

To better understand whether chromatin decondensation is a gene-independent way to prevent retina degeneration, we have investigated whether other inhibitors of chromatin condensation can prevent photoreceptor degeneration in the rd10 mouse model. We tested inhibitors for G9A/GLP, that catalyzes methylation of H3K9, and for EZH2, that catalyzes trimethylation of H3K27, and compared them to the actions of LSD1 and HDAC inhibitors. We showed that all these inhibitors that decondense chromatin are preserving retina from degeneration in mouse model of RP, rd10. The combined results provide robust support for our hypothesis that inhibition of chromatin condensation can be sufficient to prevent rod death in rd10 mice.

It is clear that the less compact chromatin has major changes in gene expression. We observed strong upregulation of rod genes involved in visual transduction in the whole retina RNA-seq, but not in the isolated rods. This strongly suggests that the observed changes are a consequence of changing numbers of rods. A question that arises is whether all of the inhibitors lead to up- or down-regulation of a few key genes, or whether neuroprotection is a broad phenomenon that can be achieved in multiple ways. To address this, we carried out RNA-seq studies for each inhibitor treatment and compared DEG and molecular pathways changes for all epigenetic inhibitors. Our previous hypothesis for inhibitors action proposed two mechanisms:1) epigenetic modifiers cause more open and accessible chromatin in rods, allowing expression of a neuroprotective genes and rod surviving; 2). suppression of inflammation in microglia and Muller glia. These two processes could be separated or work together as positive feet-back loop where normal photoreceptors, and inhibitor-treated photoreceptors do not produce any signaling molecules to activate inflammation.

In addition to their effects on visual phototransduction pathways and overall rod preservation, inhibitors of LSD1 and EZH2 caused an upregulation of genes from neuroprotective Ca^+2^ and Wnt pathways. Additionally, these modifiers downregulate inflammation pathways, such as Complement System, Leukocyte Extravasation Signaling, Natural Killer Cell Signaling, Neuroinflammation Signaling Pathway, Th2 Pathway, Toll-like Receptor Signaling. Since the changes in inflammation pathways were not detected in isolated treated rods, they probably arose in either microglia or Muller cells, both of which participate in inflammatory responses in the retina. At present we cannot distinguish whether the inhibitors are acting directly on microglia and Muller cells or whether they induce signals in rods that, in turn, affect the glial cells.

Some of the inhibitors may also have direct effects on inflammatory pathways. The LSD1 enzyme [10] participates in NF-kb activation and the same has been suggested for EZH2 [30, 31]. Our data demonstrate that KDM1a (LSD1) is 4^th^ top upstream regulator for UNC1999 and 6^th^ for DZNep indicating that these epigenetic modifiers inhibits or activates common pathway.

Such dual actions of some of the inhibitors could potentially make them more potent protective agents.

Both inhibitors of HDACs and GLP/ G9a demonstrate upregulation of NGF-stimulated transcription but mostly they down-regulate inflammation by inhibiting IL-12 Signaling and Production in Macrophages, Interferon alpha/beta signaling, Interferon gamma signaling, Neuroinflammation Signaling Pathway, Phagosome Formation. This suggests that Inhibitors of HDACs and GLP/ G9a mostly work through second mechanism proposed above – suppression of inflammation in microglia and Muller glia. HDAC inhibitors promote photoreceptors survival in several mouse models of RP and AMD [32–35] and there are numerous studies showing that HDAC inhibitors ameliorate inflammation [11, 12, 36, 37].

In summary, our findings indicate that multiple epigenetic modifiers can prevent degeneration of rod photoreceptors in a model of RP. They can achieve this by altering expression of protective and injurious genes in rods as well as modifying expression of inflammatory pathways in other retinal cells. These multiple mechanisms suggest that epigenetic modifiers are ideal agents to allow cells to reset homeostasis and counteract deleterious effects of mutations or other disease-causing stimuli. Such modifiers may prove to be valuable gene-independent therapeutics to target a range of multifactorial diseases in the retina, such as macular degeneration and diabetic retinopathy, as well as elsewhere in the nervous system.

## Supplementary materials

Supplemental figures 1-3, and supplemental table 1.

## Author Contributions

Conceptualization- EYP, JTT, CJB; methodology – EYP, LS, IA; software- LS, AS, IA; validation -EYP, LS; formal analysis and investigation – EYP; data curation – LS; writing—original draft preparation- EYP, CJB; writing—review and editing - all; funding acquisition – CJB, JTT. All authors have read and agreed to the published version of the manuscript.

## Data Availability

Raw counts and differential expression analysis generated during this study are available at GEO Submission **GSE295538**.

## Funding and Acknowledgments

This work was supported by RPB Stein Innovation award to CJB and STTR NIH R41 EY035180-01 to JTT and EYP. The Flow Cytometry Core (RRID:SCR_021134) and the Genome Sciences Core (RRID:SCR_021123) services and instruments used in this project were funded, in part, by the Pennsylvania State University College of Medicine via the Office of the Vice Dean of Research and Graduate Students and the Pennsylvania Department of Health using Tobacco Settlement Funds (CURE). The content is solely the responsibility of the authors and does not necessarily represent the official views of the University or College of Medicine. The Pennsylvania Department of Health specifically disclaims responsibility for any analyses, interpretations or conclusions.

## Supporting information

Popova EY preprint.pdf

