## Supplementary material for "Epigenetic modifiers to treat retinal degenerative diseases": Popova EY preprint.pdf

**Supplemental materials**

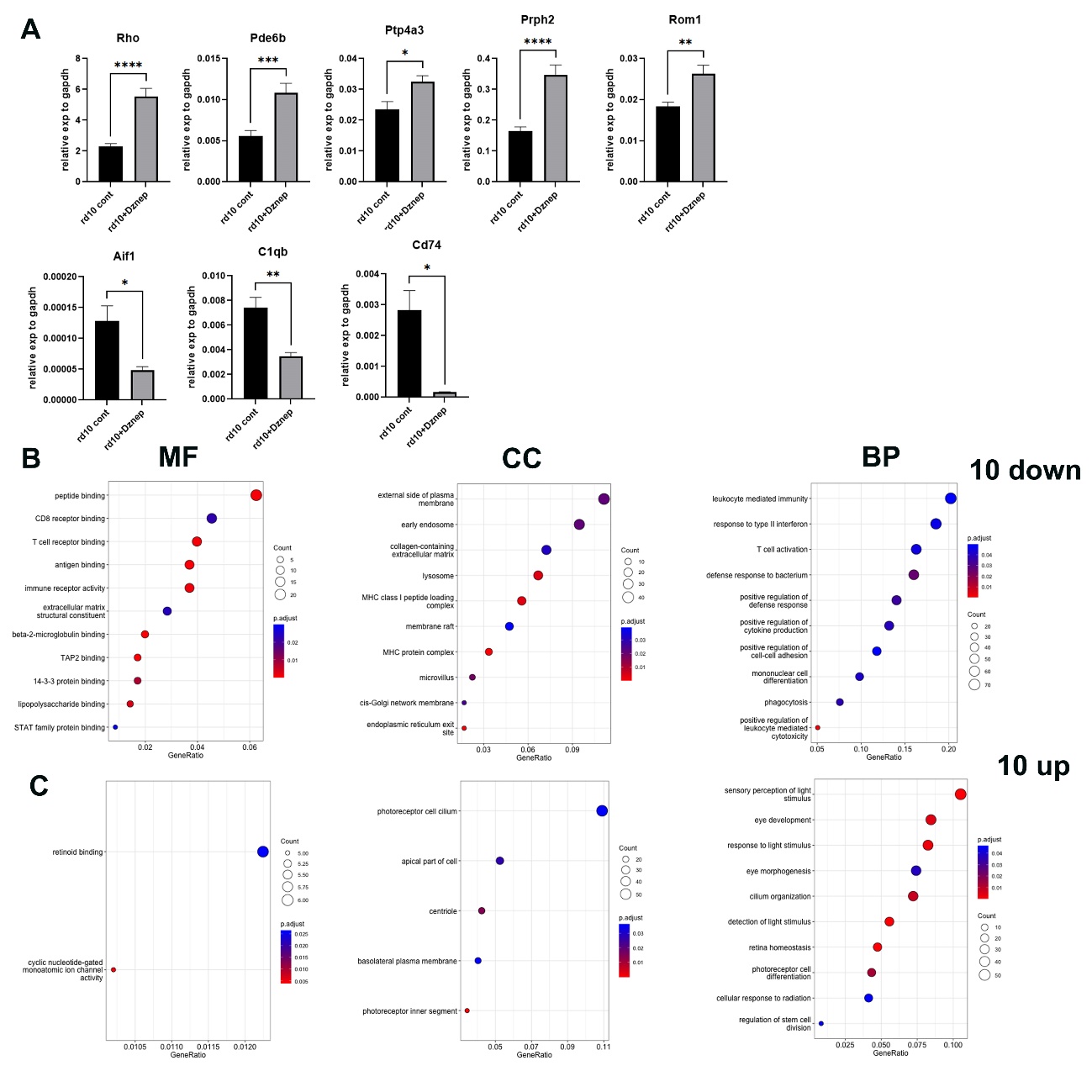

**Supplemental Fig.1. A.** Treatment of rd10 mouse model of retinitis pigmentosa with EZH2 inhibitor DZNep leads to preservation of rod photoreceptor specific genes and downregulation genes for inflammation. Comparison of rod specific gene expression levels in mice PN24 retina of rd10 mice, treated with saline, or in rd10 treated with DZNep from PN9 till PN24. Experiments were done for 2-4 biological and 3 technical replicas. The relative expression level for each gene was calculated by the 2-∆∆Ct method and normalized to GAPDH, * p<0.05; ** p<0.01; *** p<0.001, ****p<0.0001. **B-C**. Gene Ontology (GO)functional enrichment analyses of DEG between control and treated with DZNep rd10 mice retinas. The ten most significant down(**B**) and up(**C**)- regulated GO categories for Molecular Function (MF), Cell Component (CC) and Biological Process (BP).

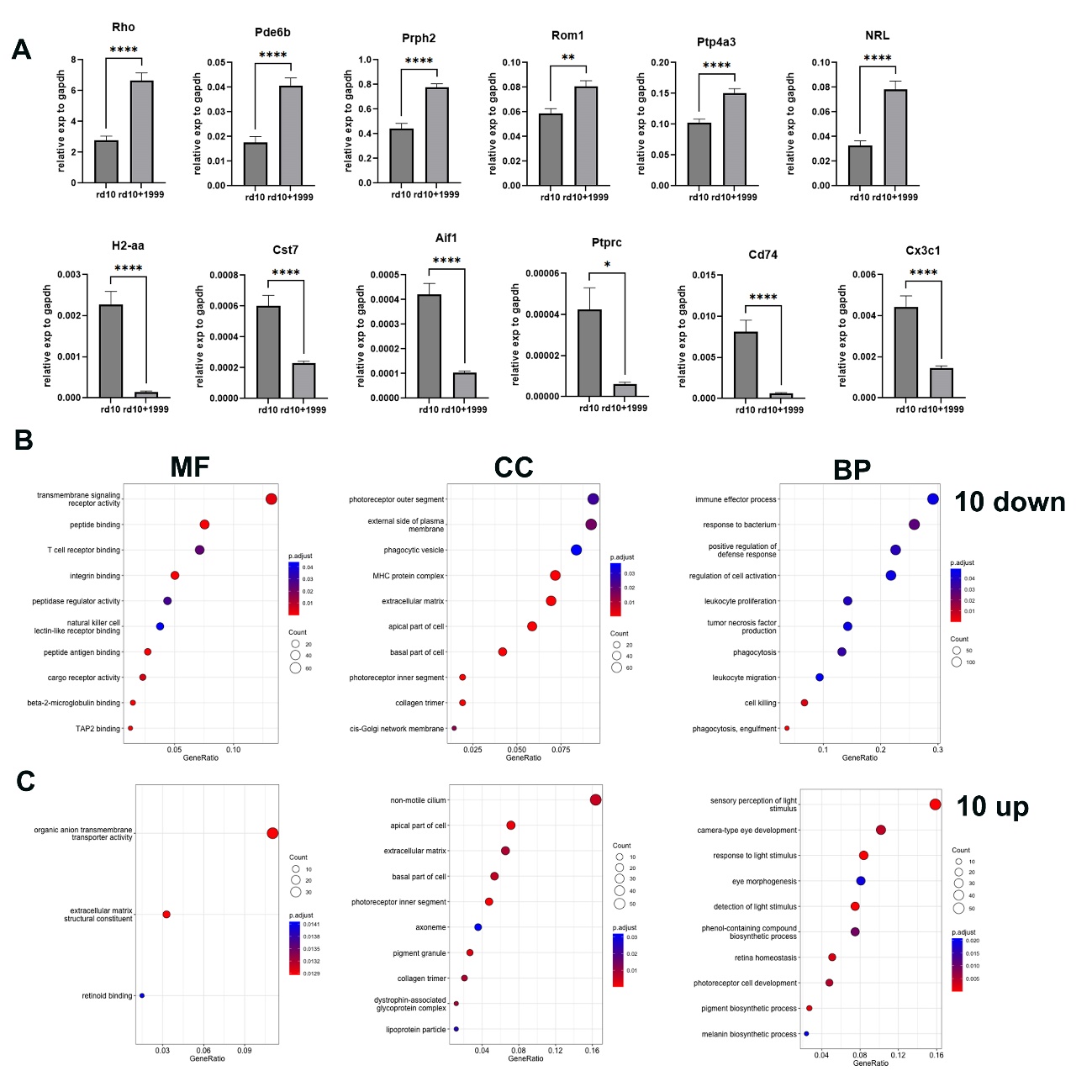

**Supplemental Fig.2. A.** Treatment of rd10 mouse model of retinitis pigmentosa with EZH2 inhibitor UNC1999 leads to preservation of rod photoreceptor specific genes and downregulation genes for inflammation. Comparison of rod specific gene expression levels in mice PN24 retina of rd10 mice, treated with saline, or in rd10 treated with UNC1999 from PN9 till PN24. Experiments were done for 3 biological and 3 technical replicas. The relative expression level for each gene was calculated by the 2-∆∆Ct method and normalized to GAPDH, * p<0.05; ** p<0.01; *** p<0.001, ****p<0.0001. **B-C**. Gene Ontology (GO)functional enrichment analyses of DEG between control and treated with UNC1999 rd10 mice retinas. The ten most significant down(**B**) and up(**C**)- regulated GO categories for Molecular Function (MF), Cell Component (CC) and Biological Process (BP).

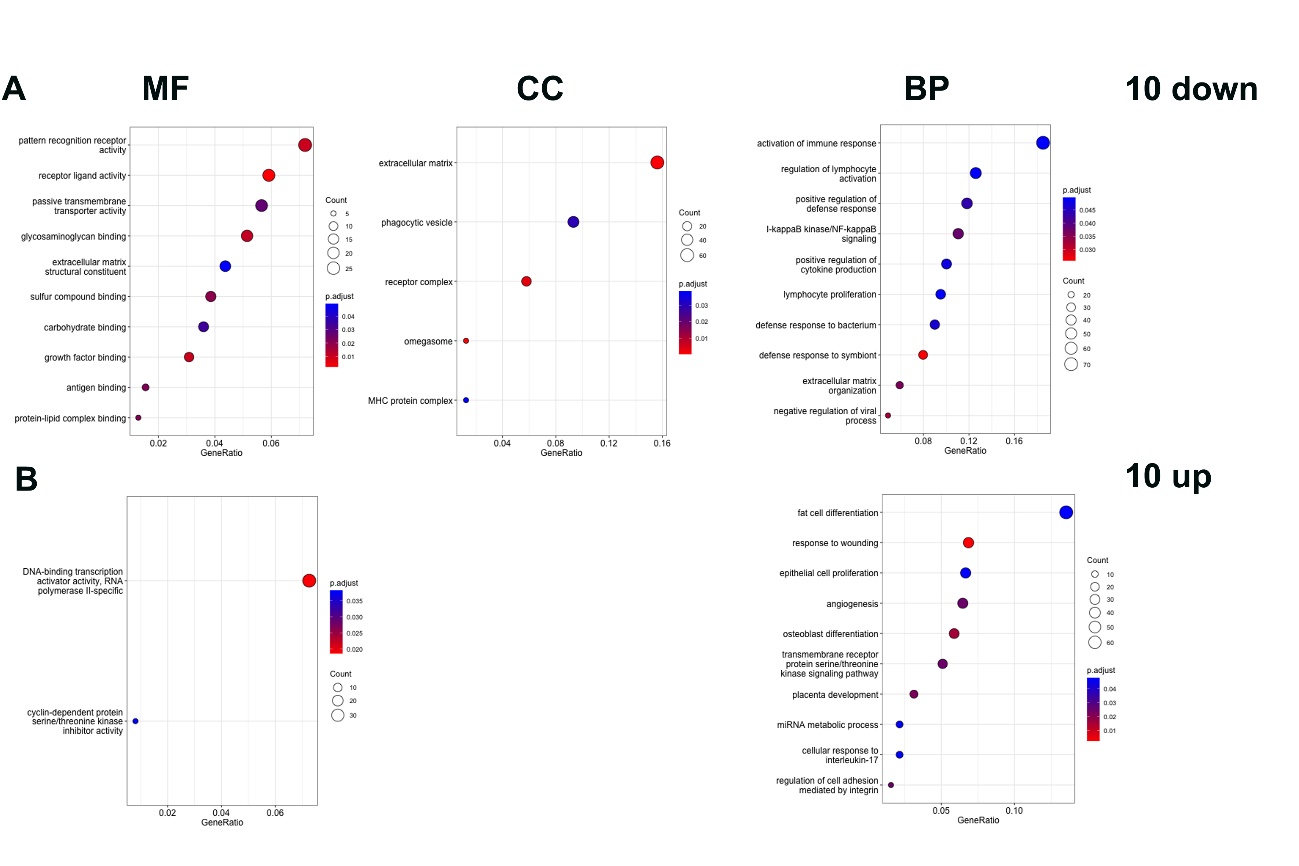

**Supplemental Fig.3.** Gene Ontology functional enrichment analyses of DEG between control and treated with Romidepsin rd10 mice retinas. The ten most significant down(**A**) and up(**B**)- regulated GO categories for Molecular Function (MF), Cell Component (CC) and Biological Process (BP).

**Supplemental table 1:**

| **Gene** | **Forward** | **Reverse** |
| --- | --- | --- |
| Aif1 | CTCCGAGGAGACGTTCAGC | TTCTCCTCATACATCAGAATCATTCTC |
| C1qb | GCTGATGAAGACACAGTGGG | GCTGTTGATGGTCCTCAGG |
| Cd74 | CAAACCTGTGAGCCAGATGC | GGTCCTGGGTCATGTTGC |
| Cst7 | CGAACTACATGCAGGAAGACC | CACTGGCAGAGGAGAACAGG |
| Cx3cr1 | CAGCATCGACCGGTACCTT | GCTGCACTGTCCGGTTGTT |
| Gapdh | Qiagen QT01658692 |  |
| H2-Aa | GTCTTGACTAAGAGGTCAAATTCC | TTCTGAGCCATGTGATGTTG |
| Nrl | GTGCCTCCTTCACCCACCTTCAGTGA | GCGTGCGGCGCCTCTGCTTCAGCCG |
| Pde6b | CTG ACG AGT ATG AGG CCA AAG | TAG GCA GAG TCC GTA TGC AGT |
| Prph2 | TGGATCAGCAATCGCTACCT | CTGTAGTAATTCAGCAGAGC |
| Ptp4a3 | TACAGAGCTTCCTCCAAGGAAA | CACGGTGTTGGGAACGG |
| Ptprc | TGCCTCACCTACACACACC | ACATGAGTCATTAGACACACTGATG |
| Rho | CTT CTC CAA CGT CAC AGG CGT | GGACCACAGGGCGATTTCAC |
| Rom1 | Qiagen #QT00172165 |  |
